# Distinct effects of different metabolic stress models on human-derived neuronal networks

**DOI:** 10.64898/2026.06.09.731175

**Authors:** Linda Collo, Eva J.H.F. Voogd, Giulia Parodi, Marloes R. Levers, Michela Chiappalone, Sergio Martinoia, Jeannette Hoffmejer, Monica Frega

## Abstract

Different *in vitro* models are widely used as experimental platforms to assess neuronal responses to metabolic stress and test potential treatments for patients with ischemic stroke. Results of those studies depend on the stress models used, and the link between cell viability–based readouts and electrophysiological activity remains poorly explored. We investigated the neuronal network activity of human-derived neuronal networks generated from human induced pluripotent stem cells (hiPSCs) under three commonly used metabolic stress models: hypoxia alone, oxygen and glucose deprivation (OGD), and hypoxia combined with different concentrations of glutamate. We aim to clarify the differences between three commonly used *in vitro* models, including the relation between microscopic and electrophysiological readouts. These conditions produced distinct effects on neuronal network activity. Hypoxia alone induced a progressive decline in activity over time. In contrast, OGD triggered a biphasic response, characterized by an early increase in activity followed by a decline. High concentration glutamate exposure under hypoxia also altered network dynamics, inducing a triphasic pattern consisting of a rapid activity decrease, a transient increase, and a subsequent decline. Across all these pathological conditions, neuronal activity progressively declined and converged toward network failure after prolonged hypoxia. Following reoxygenation, recovery was limited and condition-dependent: hypoxia alone, OGD, and high glutamate conditions showed limited recovery. On the other hand, low glutamate concentration was associated with good recovery. Microscopic assessment revealed that cellular viability was differentially affected across conditions. OGD was associated with the highest levels of cell death, whereas glutamate exposure, particularly at high concentrations, led to a marked reduction in synaptic puncta despite partial preservation of cell viability.

These findings highlight that commonly used *in vitro* ischemia models induce distinct neuronal responses and highlight the importance of integrating electrophysiological and structural analyses to better characterize metabolic stress in human neuronal networks better.

## 1. Introduction

Among the organs of the human body, the brain is the most metabolically demanding. Although it represents only 2% of body mass, it consumes 20% of oxygen and 25% of the total glucose available in the body^1^, making it highly vulnerable to interruptions in blood flow. Indeed, acute ischemic stroke, which is caused by cerebral blood flow disruption, is a leading cause of mortality and long-term disability ^2,3^. Reduced perfusion limits oxygen and glucose delivery, triggering a cascade of events including energy failure, excitotoxicity, oxidative stress, inflammation, and ultimately neuronal death ^4,5^.

Current treatments rely on acute recanalization via thrombolysis or endovascular thrombectomy ^6^. However, strict eligibility criteria and limited therapeutic temporal windows restrict their use, and many patients still retain significant neurological deficits despite successful reperfusion ^7–9^. Therefore, additional neuroprotective strategies are needed to prevent secondary injury and improve functional recovery.

In this context, *in vitr*o models provide a controlled and reproducible platform to study ischemic mechanisms and evaluate potential therapeutic approaches.

Three main *in vitro* models are commonly used to study neuronal ischemic injury. Hypoxia-based models reduce oxygen availability to mimic the conditions of ischemic penumbra ^10^. Oxygen–glucose deprivation (OGD), which involves the simultaneous removal of oxygen and glucose, reproduces the metabolic and excitotoxic stress characteristic of the first phase of ischemia ^11,12^. Excitotoxicity-based models expose cells to excessive glutamate, mimicking the later stages of the ischemic cascade ^13^. Most *in vitro* ischemia models have been developed using rodent cells ^14–19^: while these systems are widely employed, rodent neurons differ from human neurons in gene expression, receptor composition, and cellular responses to ischemic stress, which can limit the translational relevance of the findings ^20^. To overcome these limitations, recent studies have shifted toward human-based systems. Pires Monteiro et al. (2021) developed a human induced pluripotent stem cells (hiPSCs)-based model of the ischemic penumbra using hypoxia ^10^. Subsequent studies have combined OGD with hiPSCs-derived neurons ^21^. Similarly, human iPSCs-derived neurons have been used to study glutamate-induced excitotoxicity, allowing investigation of calcium dysregulation, receptor-mediated responses (AMPA/NMDA) ^22,23^.

Although hypoxia-induced effects have been investigated in both rodent and human *in vitro* models of ischemic injury using electrophysiological approaches ^10,17^, widely used paradigms such as OGD and glutamate-induced excitotoxicity are still predominantly assessed through viability-based readouts, focusing on endpoints such as cell apoptosis and structural damage. Nevertheless, electrophysiological analyses provide complementary information by monitoring neuronal activity over time, including transitions from reversible to irreversible injury. Furthermore, combining electrophysiological measurements with molecular analyses enables a more comprehensive understanding of the mechanisms underlying functional changes, supporting the evaluation of novel treatments.

Although (1) hypoxia, (2) OGD, and (3) glutamate-induced excitotoxicity are well-established *in vitro* models of ischemic injury, a direct and systematic comparison of their effects in human neuronal cultures is still lacking. Here, we compare these three commonly used *in vitro* ischemia models, by combining electrophysiological recordings with complementary analyses of cell viability, apoptosis, and synaptic density.

Overall, our results show that different paradigms yield distinct temporal dynamics but converge on a similar functional impairment at the network level, while the underlying cellular and synaptic alterations differ substantially. These findings highlight that commonly used *in vitro* models can generate distinct but partially overlapping responses, emphasizing the importance of carefully selecting and interpreting the experimental paradigm when studying ischemia-related neuronal dysfunction.

## 2. Materials and Methods

### 2.1. HiPSCs generation

Two previously characterized hiPSCs lines were genetically modified to generate homogeneous population of excitatory and inhibitory neurons by forced overexpression of Neurogenin 2 (*Ngn2*) and Achaete-scute homolog 1 (*Ascl1*) transcription factors, respectively. The hiPSCs lines were received in frozen vials from Prof. N. Nadif Kasri (Radboud University, Nijmegen, The Netherlands) and were originally derived from fibroblasts. Line 1 (healthy 30-year-old female) was reprogrammed via episomal reprogramming (Coriell Institute for medical research, GM25256). Line 2 (healthy 51 years-old male) was reprogrammed via a non-integrating Sendai virus (KULSTEM iPSC core facility Leuven, Belgium, KSF-16-025). Cultures were maintained in E8Flex medium (Thermo Fisher Scientific) supplemented with E8 supplements (50X, Thermo Fisher Scientific), G148 (50 μg/ml, Sigma Aldrich) and puromycin (0.5 μg/ml, Sigma Aldrich). The cultures were maintained at 37°C and 5% CO_2_ in the incubator. The medium was refreshed every two days.

### 2.2. HiPSCs differentiation

Following the protocol presented by Mossink et al. (2022), at Day *In Vitro* (DIV) 0, Ngn2- and Ascl1-positive hiPSCs were detached with Accutase and plated together in a proportion equal to 80:20 (E:I) on 24-well MEAs (Multi-channel System, for electrophysiological characterization) or glass coverslips into a sterile 24-well (for immunocytochemical staining) (Fig. 1a) ^24^. Devices were pre-coated with poly-l-ornithine (50 μg/ml, 5 hours, Sigma Aldrich) and then with human laminin (20 μg/ml, overnight, Bioconnect). Doxycycline (4 µg/ml, Sigma Aldrich) and forskolin (4 µg/ml, Thermo Fisher Scientific) were added to induce overexpression of *Ngn2* and *Ascl1*, thereby supporting differentiation into excitatory and inhibitory neurons, respectively. To further support maturation, new-born (P1) Wistar rat cortical astrocytes were added to the neuronal networks in a 1:1 ratio two days after plating (in agreement to Dutch and European laws and the guidelines of the Dutch Animal Use Committee) (Fig. 1a). The cultures were maintained in Neurobasal medium supplemented with B27 supplements (2%, Gibco), primocin (0.1 μg/ml, Invivogen), GlutaMax 100X (1%, Thermo Fisher Scientific), human Brain-Derived Neurotrophic Factor (10 ng/ml, Stemcell Technologies), human Neurotrophin-3 (10 ng/ml, Stemcell Technologies), doxycycline (4 µg/ml), and forskolin (4 µg/ml). Cytosine β-D-arabinofuranoside (Ara-C, 2 μM, Sigma Aldrich) was added only in the medium on the day after the addition of the astrocytes to block any proliferating cells. The medium was additionally supplemented with 2.5% FBS (Sigma Aldrich) to support astrocytes’ viability from DIV10 onward. After 3 weeks of treatment with doxycycline and forskolin, the cells were considered fully differentiated into neurons and both compounds were removed from the medium. In this experimental protocol, the medium was refreshed every 2 days. Cells were kept in an incubator in stable conditions (37 °C, 100% humidity, and 5% of CO_2_) until the day of the experiment (DIV49).

**Figure 1.**
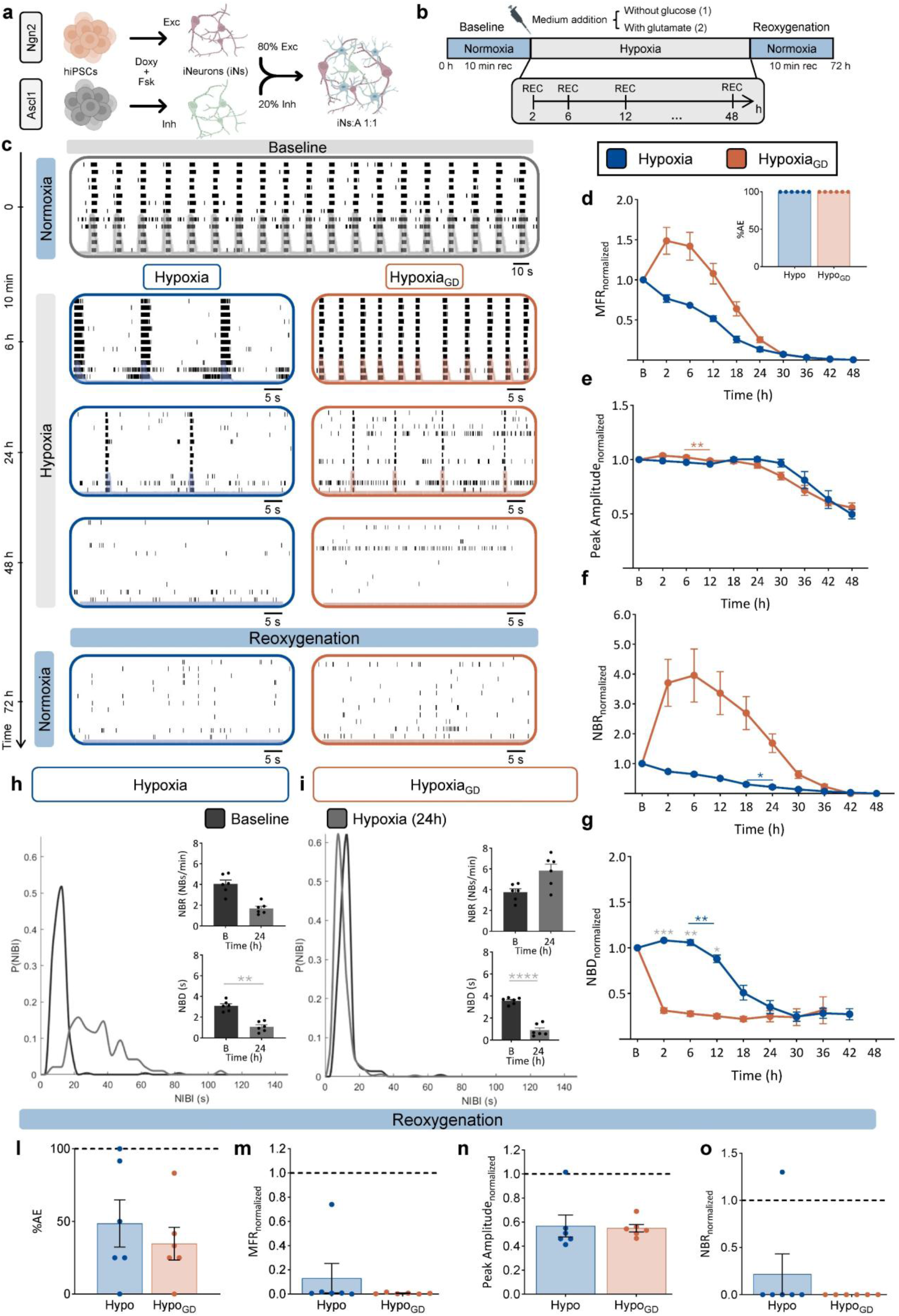
Effect of oxygen and glucose deprivation (OGD) on neuronal network activity. **a)** Sketch of the neuronal differentiation protocol: hiPSCs were differentiated into excitatory (E) and inhibitory (I) iNeurons (iNs) with doxycycline and forskolin treatment. Neurons were co-plated in a ratio equal to 80:20 (E: I). Rat astrocytes (A) were added at a 1:1 ratio (iNs:A) to support neuronal growth. **b)** Schematic representation of the performed experimental protocol with different conditions: only hypoxia, oxygen glucose deprivation, and hypoxia in combination with glutamate. After 10-minute recordings in normoxia condition (37°C, 20% O2, 75% N_2_), the culture medium was changed and the neuronal cultures were subjected to 48 hours of hypoxia (37°C, 2% O_2_, 93% N_2_), in which the activity was recorded for 10 minutes after 2 hours and then 10 minutes every 6 hours. The last phase of the experimental protocol consisted of 24 hours of normoxia (i.e. reoxygenation, 37°C, 20% O_2_, 75% N_2_, for a total of 72 hours of experiment), in which the activity was recorded for 10 minutes. **c)** *<u>Ontop:</u>* representative raster plot showing five minutes of electrophysiological activity during normoxia (i.e. baseline). *<u>Inthemiddle:</u>* representative raster plots during hypoxia phase showing one minute of electrophysiological activity for hypoxia (in blue) and hypoxia_GD_ (in orange) after 6, 24, and 48 hours of oxygen deprivation. *<u>Onthebottom:</u>* representative raster plots showing one minute of electrophysiological activity during the reoxygenation phase for hypoxia (in blue) and hypoxia_GD_ (in orange). Black dots represent detected spikes, overlapped the cumulative instantaneous firing rate profile. **d-g)** Graphs showing the profile during hypoxia (in blue) and hypoxia_GD_ (in orange) of: normalized firing rate, *<u>inset:</u>* bar graphs showing percentage of active electrodes**(d)**; normalized peak amplitude **(e)**; normalized network bursting rate **(f)**; normalized network burst duration **(g)**. Data are shown with the mean (dots) and standard error of the mean (whiskers). **(h-i)** NIBI distributions in networks exposed to hypoxia **(h)** and hypoxia_GD_ **(i)** during baseline (black) and after 24 hours of hypoxia (grey). *<u>Inset:</u>* bar plots of network bursting rate and network burst duration during baseline and after 24 hours of hypoxia. **(l-o)** Bar graphs showing: percentage of active electrodes **(l)**; normalized firing rate **(m)**; normalized peak amplitude **(n)**; normalized network bursting rate **(o)** of hypoxia (in blue) and hypoxia_GD_ (in orange) after 24 hours of reoxygenation. The dashed line represents the baseline value in normoxia. In the bar plots, data are shown with the mean (bar), single values (dots), and standard error of the mean (whiskers). In figure d-g asterisks with bars underneath indicate comparisons between consecutive time points within the same group, whereas grey asterisks denote comparisons between the two groups at the same time point *p < 0.05, **p < 0.01, ***p < 0.001, ****p < 0.0001. Data were obtained from n = 6 independent neuronal cultures for hypoxia and n = 6 independent neuronal cultures for hypoxia_GD_.

### 2.3. Experimental protocol

To model ischemic stroke *in vitro*, the hiPSCs-derived neuronal networks were exposed to (1) hypoxia, (2) OGD (hypoxia_GD_) or (3) hypoxia in combination with different concentrations of glutamate (100, 200, and 500 µM; hypoxia_100µM_,hypoxia_200µM_,hypoxia_500µM_, respectively). Each glutamate concentration was tested in a separate well, meaning that individual wells were exposed to only one glutamate dose throughout the experiment.

All the experiments were performed at DIV 49 in a computer-controlled climate chamber. As reported in Figure 1b, the experiment consisted of three different phases. Neuronal networks were initially exposed to normoxia (20% O_2_, 75% N_2_, 5% CO_2_, “Baseline”). Subsequently, cells were subjected to the three different conditions for 48 hours: (1) hypoxia (2% O_2_, 93% N_2_, 5% CO_2_), (2) hypoxia_GD_, achieved by inducing hypoxia and replacing the culture medium with glucose-free medium, and (3) glutamate exposure, in which hypoxia was combined with different concentrations of glutamate (Fig. 1b). After hypoxia, the networks were returned to normoxia for 24 hours (Fig. 1b, “Reoxygenation”) and the wells were filled with fresh culture medium containing glucose. The effect on the neuronal networks was evaluated with both electrophysiological recordings and immunocytochemistry.

### 2.4. Electrophysiological recordings and data analysis

The electrophysiological recordings were performed using Multiwell-MEA-System (Multi Channel Systems, Reutlingen, Germany) through 24 wells Micro-Electrode Arrays (MEAs). Each well contained 12 embedded electrodes (4 × 4 grid) with a 30 µm diameter and 200 µm pitch. All recordings were acquired at a sampling rate of 10 kHz for 10 minutes. After an acclimatization period, baseline activity was recorded under normoxic conditions. During the hypoxic phase, neuronal activity was recorded after 2 hours and subsequently every 6 hours (Fig. 1b). Finally, the activity was recorded after 24 hours of normoxia (Fig. 1b).

During the recordings, the signals were filtered using a low-pass filter (4th order Butterworth filter, 3.5 kHz) and a high-pass filter (2nd order Butterworth filter, 100 Hz). The data analysis was performed using in-house code implemented in MATLAB (The MathWorks, Natick, MA, USA) and SpyCode ^25^. Briefly, spike detection was carried out with the Precision Time Spike Detection (PTSD) algorithm ^26^. The noise threshold for detecting individual spikes was set at 10 times the standard deviation of the baseline noise. The peak lifetime period, corresponding to the spike duration, and the refractory period, corresponding to the minimum interval between consecutive spikes, were both set to 2 ms. Electrodes exhibiting a mean firing rate (MFR; defined as the number of spikes per unit time) over 0.1 spikes/s (spk/s) were considered for analysis ^27^. Firing activity decay of the glutamate-treated group was modelled using a mono-exponential function y = *e*^*bt*^, where b represents the decay constant. The fitting was performed in MATLAB using a non-linear regression analysis. Subsequently, the peak amplitude was defined as the difference between the initial peak and the subsequent opposite peak.

The burst detection was performed based on the logarithmic ISI distribution ^28^. An event was considered a burst if composed of at least 10 spikes. Electrodes with a mean bursting rate (MBR, the number of bursts per unit of time) greater than 0.4 bursts/min were considered as bursting channels ^27^. Then, an event was defined as a network burst if 30% of the bursting channels simultaneously presented activity. The network bursting rate (NBR, the average frequency of synchronized burst event) was computed together with the network burst duration (NBD). The MFR, NBR and NBD of each well were extracted from each time point and normalized with respect to the MFR, NBR, and NBD in normoxia (baseline) of the same well.

Finally, we analyzed the distributions of network inter-burst intervals (NIBI) during the baseline phase in normoxia, after 24 hours of hypoxia, and after 72 hours (24 hours of normoxia). For each analysed group, the baseline of that same group was used as a reference.

### 2.5. Live/Dead Assay (LDA)

The viability of neuronal networks cultured on glass coverslips was assessed using a live/dead assay in four conditions: normoxia, hypoxia, hypoxia_GD_, and hypoxia_500µM_. Apoptotic cells were defined as cells undergoing programmed cell death, whereas dead cells were defined as cells exhibiting loss of membrane integrity, independently of the cell death pathway. CellEvent (1:500, ThermoFisher Scientific) was added to each well and incubated for 30 minutes at 37°C to stain apoptotic cells (green). After that, the hypoxic period started for 48 h. At the end of the hypoxic period, propidium Iodide (PI, 1:1000, Invitrogen) was added to each well and incubated for 15 minutes at 37°C to detect dead cells (red). Following incubation, the coverslips were rinsed with phosphate-buffered saline (PBS) and fixed with 3.7% paraformaldehyde (PFA, Sigma Aldrich) for 15 minutes at room temperature. After washing with PBS, DAPI (1:1000, Sigma Aldrich) was added and incubated for 10 minutes at room temperature to stain cell nuclei (blue). Coverslips were washed again with PBS and mounted onto glass microscope slides using Mowiol (Sigma Aldrich). The slides were maintained in the dark over/night at room temperature and subsequently stored at 4°C.

Neuronal networks have been included in the dataset only when they showed good quality (i.e. cell density allowing for proper neuron-electrode coupling and even distribution of cells. Images of the cultures were acquired at 40× magnification (0.085 μm/pixel) using a Nikon Eclipse 50i epifluorescence microscope (Nikon, Japan). Pre-processing of images was performed using a custom MATLAB script (The MathWorks, Natick, MA, USA). The quantification of apoptotic and dead cells was conducted manually with ImageJ. Dead cells were defined as those positive for PI (red) or for both CellEvent and PI (yellow, resulting from the colocalization of green and red signals). Apoptotic cells were defined as those positive for CellEvent only (green). The number of living cells was calculated by subtracting the sum of apoptotic and dead cells from the total number of DAPI-positive cells (blue).

### 2.6. Quantification of synaptic puncta

After exposure to normoxia, hypoxia, hypoxia_GD_, and hypoxia_500µM_, hiPSCs-derived neuronal networks were fixed with 3.7% paraformaldehyde (PFA, Sigma Aldrich) for 15 minutes at room temperature (RT). Following fixation, the networks were washed with PBS and permeabilized for 5 minutes in 0.2% Triton X-100 (Sigma Aldrich) in PBS. Non-specific bindings were blocked by incubating the cultures in 2% bovine serum albumin (BSA) for 30 minutes at RT.

Primary antibody staining was performed overnight at 4°C using rabbit anti-Bassoon (1:500) and mouse anti-PSD-95 (1:50). The coverslips were then washed with PBS and incubated for 1 hour at RT with secondary antibodies: goat anti-rabbit AF568 (1:2000, A11036, Invitrogen) and goat anti-mouse AF488 (1:2000, A11029, Invitrogen), followed by PBS washes. Nuclei were counterstained with DAPI (1:1000, Sigma Aldrich) for 10 minutes at RT. Finally, coverslips were washed with PBS, mounted with Mowiol (Sigma Aldrich), and allowed to dry overnight.

Neuronal networks have been included in the dataset only when they showed good quality (i.e. cell density allowing for proper neuron-electrode coupling and even distribution of cells. Images were acquired at 60× magnification (0.057 μm/pixel) using a Nikon Eclipse 50i epifluorescence microscope (Nikon, Japan. Synaptic puncta were quantified using Synapse Counter, an ImageJ plug-in developed for rapid, automatic, and unbiased quantification of synaptic marker proteins ^29^. Synapse Counter allows for the measurement of both the number and size of presynaptic and postsynaptic clusters, as well as the identification of co-localizing puncta, indicative of well-formed and mature synapses. The size thresholds for presynaptic and postsynaptic particles were set in pixels^2^ (or voxels).

### 2.7. Statistical analysis

First the evaluation of outliers was performed with an in-house code implemented in MATLAB (The MathWorks, Natick, MA, USA) using a median-based approach, in which values exceeding three scaled median absolute deviations from the median were considered outliers. Then, statistical analysis was performed using Prism (GraphPad Software Inc.). Data normality was assessed using the Shapiro–Wilk test. Depending on data distribution and experimental design, parametric or non-parametric tests were applied. Comparisons involving multiple time points were analyzed using mixed-effects models, with Sidak’s multiple comparison test. Non-normally distributed data were analyzed using the Kruskal–Wallis test with Dunn’s multiple comparisons test. For data that were normally distributed Brown–Forsythe and Welch’s ANOVA test was applied with Dunnet multiple comparison test. For the comparison between the two groups, a Welch test or a Kolmogorov-Smirnov were applied. Statistical significance was set at *p* < 0.05.

## 3.. Results

### 3.1. Hypoxia and glucose deprivation induce transient hyperactivity followed by network failure

We differentiated hiPSCs into neuronal networks and investigated the effect of 48 hours of hypoxia alone (hypoxia) and hypoxia combined with glucose deprivation (hypoxia_GD_) on neuronal activity (Fig. 1a, b). By seven weeks *in vitro*, neuronal networks grown on MEAs on physiological oxygen conditions (normoxia) were functionally mature, exhibiting typical stable electrophysiological activity composed by spikes and synchronous bursts (Fig. c, “Normoxia”).

Hypoxia induced a progressive decline in activity over time, characterized by a gradual reduction in firing rate, network burst frequency and duration, and number of active electrodes, consistent with previous reports (Fig. 1c-f, blue) ^10^. In contrast, hypoxia_GD_ elicited a biphasic response, with an initial increase in activity, peaking at 6 hours, followed by a progressive decline (Fig. 1c-f, orange). During the first 24 hours, hypoxia_GD_ networks displayed higher firing rates and more frequent but shorter bursts compared to both normoxia and hypoxia (Fig. 1d, f-g). Over time, both conditions converged toward severe functional impairment. By 48 hours, neuronal activity was largely abolished in both models, with only sparse, uncoordinated spiking and no detectable network bursts. Analysis of network inter-burst interval (NIBI) distributions further supported these dynamics (Fig. 1h-i): under normoxic conditions (baseline), the distribution showed a sharp, unimodal peak, reflecting regular network bursting activity. Under hypoxia, the distribution progressively became broader and irregular, consistent with gradual network disorganization. In contrast to hypoxia, hypoxia_GD_ showed an early shift toward shorter network inter-burst intervals, followed by complete loss of network bursting activity, with no recovery after reoxygenation (Fig. 1h-i).

After 48 hours of hypoxia, neuronal networks were returned to normoxic conditions to assess recovery. Following reoxygenation, recovery was minimal in both models (Fig. 1c, l–o, “Reoxygenation”).

### 3.2. Glutamate modulates neuronal activity under hypoxia in a concentration-dependent manner

We subsequently investigated the combined effects of hypoxia and glutamate at different concentrations (i.e., 100 µM, 200 µM, and 500 µM) on the dynamics of hiPSCs-derived neuronal networks, comparing their effects relative to hypoxia alone. The raster plots revealed a decline in neuronal activity under glutamate exposure, characterized by faster and distinct temporal dynamics compared to hypoxia alone, with a more pronounced and concentration-dependent effect at higher concentrations (Fig. 2a).

**Figure 2.**
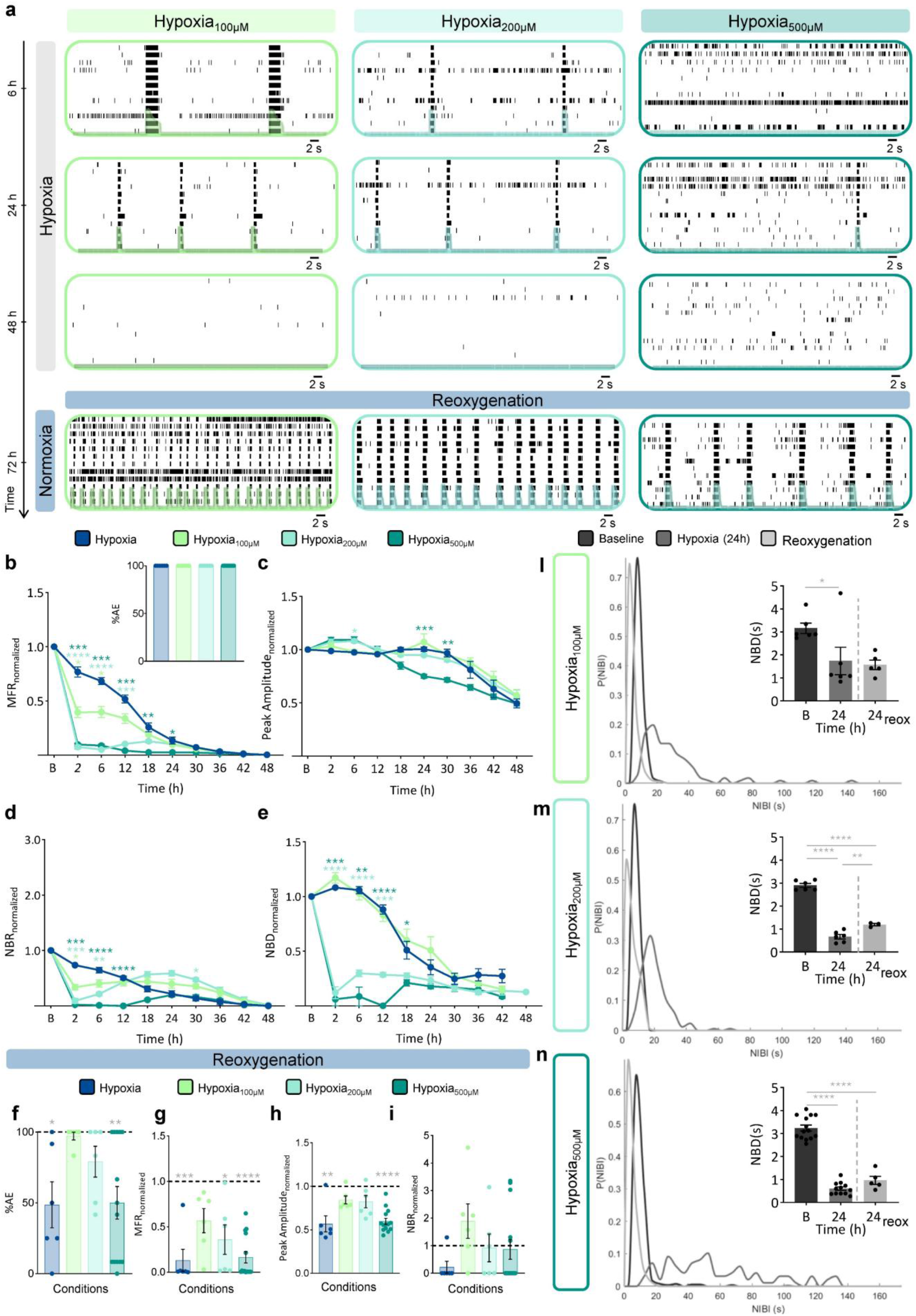
Effect of hypoxia and glutamate on neuronal network functionality. **a)** *<u>Ontop:</u>* representative raster plots showing one minute of electrophysiological activity of neuronal networks under hypoxia_100µM_ (light green), hypoxia_200µM_ (mint green), and hypoxia_500µM_ (dark teal) after 6, 24, and 48 hours of hypoxia. *<u>Onthebottom:</u>* representative raster plots showing one minute of electrophysiological activity during the reoxygenation phase (i.e. 24 hours normoxia) for hypoxia_100µM_ (light green), hypoxia_200µM_ (mint green), and hypoxia_500µM_ (dark teal). Black dots represent detected spikes, overlapped the cumulative instantaneous firing rate profile. **b-e)** Graphs showing the profile during hypoxia (in blue), hypoxia_100µM_ (light green), hypoxia_200µM_ (mint green), and hypoxia_500µM_ (dark teal) of: normalized firing rate (*<u>inset:</u>* bar graphs showing percentage of active electrodes) **(b)**; normalized peak amplitude **(c)**; normalized network bursting rate **(d)**; normalized network burst duration **(e)**. Data are shown with the mean (dots) and standard error of the mean (whiskers). **(f-i)** Bar graphs showing: percentage of active electrodes **(f)**; normalized firing rate **(g)**; normalized peak amplitude **(h)**; normalized network bursting rate **(i)** after 24 hours of reoxygenation. The dashed line represents the baseline value in normoxia. **(l-n)** NIBI distributions in networks exposed to hypoxia_100µM_ **(l)**, hypoxia_200µM_ **(m)** and hypoxia_500µM_ **(n)** during baseline (black), after 24 hours of hypoxia (grey), and 24 hours of reoxygenation (light grey). *<u>Inset:</u>* bar plots of network burst duration during baseline, after 24 hours of hypoxia, and 24 hours of reoxygenation. In the bar plots, data are shown with the mean (bar), single values (dots) and standard error of the mean (whiskers). Asterisks in figure b-e denote significant comparisons between the two groups at the same time point, while the asterisk in figure i-n denote significant comparisons between the group and the baseline .*p < 0.05, **p < 0.01, ***p < 0.001, ****p < 0.0001. Detailed statistical analyses, including the specific group comparisons and corresponding values, are provided in Supplementary Table S1, S2, S3 and S4.Data were obtained from n = 6 independent neuronal cultures for hypoxia, n = 6 independent neuronal cultures for hypoxia_100µM_, n = 6 independent neuronal cultures for hypoxia_200µM_ and n = 14 independent neuronal cultures for hypoxia_500µM_.

Overall, across all glutamate-treated conditions, from a quantitative perspective, firing activity decreased more rapidly than under hypoxia alone and approached near-zero values between 36 and 48 hours (Fig. 2a,b). Peak amplitude followed a similar trend, with earlier and more pronounced reduction at higher glutamate concentrations (Fig. 2c).

To better characterize the temporal evolution of neuronal activity, three temporal windows were defined based on the observed dynamics of the glutamate response, corresponding to first, second, and third temporal phases.

During the first temporal phase (0-6 hours), glutamate exposure induced an immediate, dose-dependent reduction in neuronal activity. This effect was reflected in faster decay kinetics compared to hypoxia (decay constant: hypoxia = -0.13; hypoxia_100µM_ = -0.46; hypoxia_200µM_ = -1.29; hypoxia_500µM_ = -1.16) (Fig. 2b). Similarly, network bursting was rapidly suppressed across all conditions and was completely abolished at the highest concentration (500 µM) (Fig. 2d). Network burst duration (NBD) was markedly reduced at higher concentrations (200 and 500 µM), whereas it remained comparable to hypoxia alone at 100 µM (Fig. 2e).

During the second temporal phase (6–24 hours), neuronal networks exposed to glutamate showed a transient, dose-dependent recovery of bursting activity, with network bursts re-emerging at reduced levels compared to baseline, whereas under hypoxia alone, activity continued to decrease. This recovery was more pronounced at lower glutamate concentrations. NBD remained relatively stable at hypoxia_100µM_ but remained reduced at higher concentrations (Fig. 2l). In contrast, mean firing rate (MFR) did not show a comparable recovery and stayed low across all conditions.

At the third temporal phase (24–48 hours), all conditions converged toward severe functional impairment. By 48 hours, network-level activity was abolished, with only sparse spiking detectable (Fig. 2a), and both MFR and NBR approached minimal values across all conditions.

Following reoxygenation, neuronal activity showed a partial and condition-dependent recovery. Hypoxia showed limited recovery, with persistently low activity levels. A similar level of impairment was observed at the highest glutamate concentration (hypoxia_500µM_). Networks exposed to 100 µM glutamate showed substantially better recovery, with several parameters returning to levels not significantly different from baseline. Intermediate concentrations (hypoxia_200µM_) showed partial recovery, remaining between these two extremes. This pattern was consistently observed across multiple measures, including the percentage of active electrodes, MFR, peak amplitude, and network bursting activity (Fig. 2f–i).

Consistent with these observations, analysis of network inter-burst interval (NIBI) distributions revealed marked alterations in network bursting regularity. During baseline (in black), all conditions exhibited sharp unimodal distributions, indicative of regular network bursting activity (Fig. 2l-n). After 24 hours of hypoxia, distributions become broadened, indicating increased variability in network burst timing. The hypoxia_500µM_ showed a more irregular and multimodal distribution, suggesting disrupted network organization (Fig. 2n). After reoxygenation, distributions shifted toward shorter network inter-burst intervals and became narrower, consistent with partial restoration of network bursting regularity (Fig. 2l-n).

### 3.3. Hypoxia, glucose deprivation and glutamate exposure differentially affect cell viability and synaptic integrity

To assess cellular viability, we evaluated the number of live, dead, and apoptotic cells in the conditions of normoxia, hypoxia, hypoxia_GD_, and hypoxia_500µM_. In parallel, we assessed the number of synaptic puncta in each experimental group. The hypoxia_500µM_ condition was selected for this analysis as it showed the strongest electrophysiological impairment.

Representative images (Fig. 3a) and quantitative analysis (Fig. 3b) showed differences in cell viability across conditions. Normoxic cultures displayed the highest proportion of live cells compared to all hypoxic groups (Fig. 3b). Under hypoxia, viability was markedly reduced and apoptotic cells were significantly increased compared to normoxia (p = 0.00013 and p = 0.00002, respectively). In contrast, hypoxia_GD_ showed a different profile, characterized by a higher proportion of dead cells rather than apoptosis. The hypoxia_500µM_ condition exhibited an intermediate phenotype, with partial preservation of viability compared to hypoxia and hypoxia_GD_.

**Figure 3.**
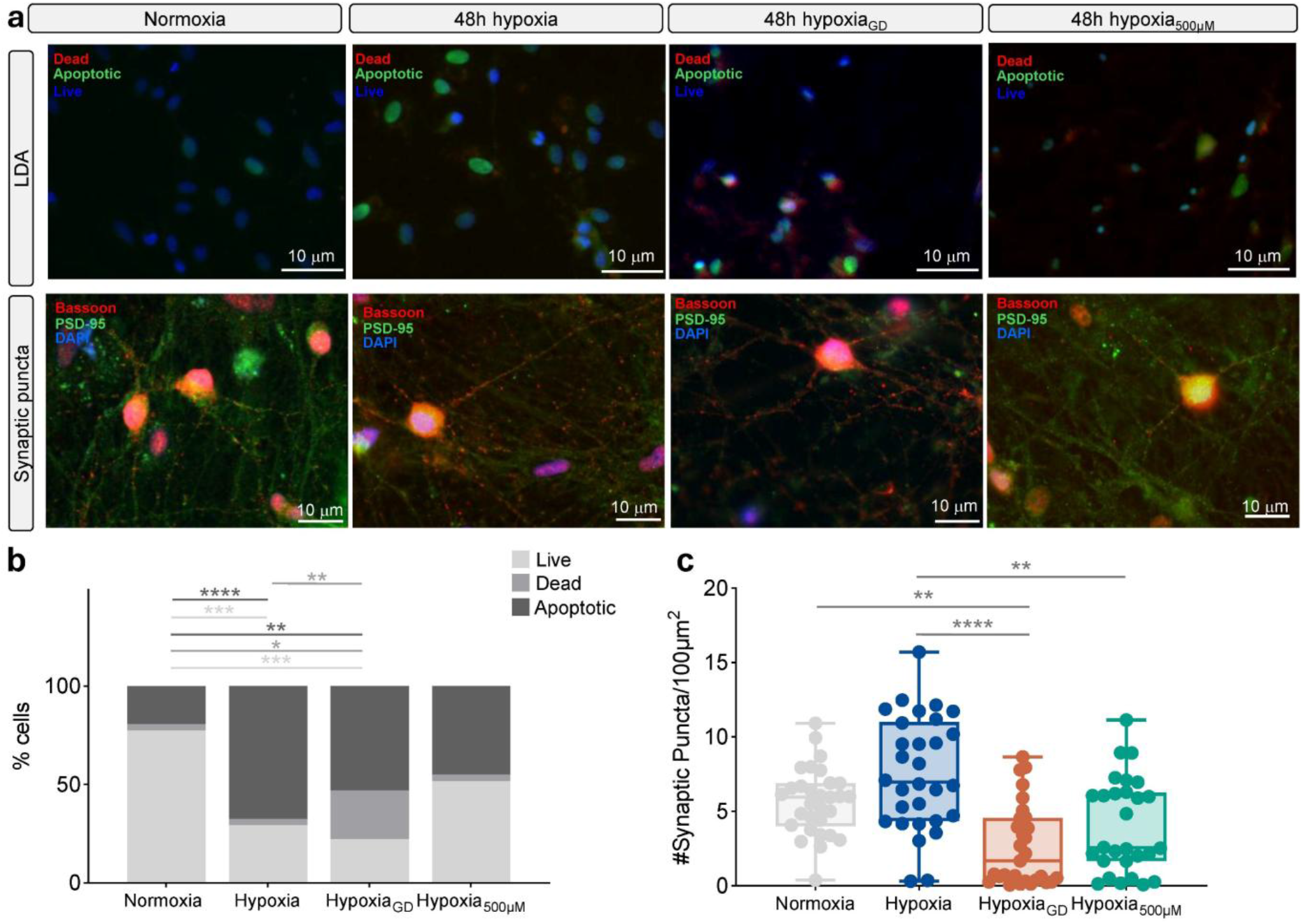
Effect of GD and glutamate combined with hypoxia on cellular viability and synaptic puncta. **a)** First row: representative images showing live (positive for DAPI; blue), dead (positive for Propidium Iodide; red) and apoptotic cells (positive for Cell Event; green) of normoxia, hypoxia, hypoxia_GD_ and hypoxia_500µM_, after 48 hours. Second row: representative images showing synaptic puncta (yellow) obtained by overlapping pre-synaptic (Basson; red) and post-synaptic (PSD-95; green) of normoxia, hypoxia, hypoxia_GD_ and hypoxia_500µM_, after 48 hours. **b)** Bar graphs showing the number of live (light grey), dead (grey), and apoptotic (black) cells in normoxia, hypoxia, hypoxia_GD_ and hypoxia_500µM_, after 48 hours of hypoxia. **c)** Box plot showing the number of synaptic puncta in normoxia (grey), hypoxia(blue), hypoxia_GD_ (orange) and hypoxia_500µM_ (dark teal), after 48 hours. In the box plots, data are shown with the median (central line), the interquartile range (box, 25th–75th percentiles), and the minimum and maximum values (whiskers). * p < 0.05, ** p < 0.01, *** p < 0.001, ****p < 0.0001. Data for LDA were obtained from 30 fields per condition. Synaptic puncta quantification was performed on n = 30 fields for normoxia, n = 30 fields for hypoxia, n = 29 fields for hypoxia_GD_, and n = 28 fields for hypoxia_500µM_.

We next examined the number of synaptic puncta (Fig. 3a, second row). While the hypoxia group maintained synaptic puncta levels comparable to normoxia, both hypoxia_500µM_ and hypoxia_GD_ groups showed a reduced number of synaptic puncta (Fig. 3a-c). Quantitative analysis confirmed these observations (Fig. 3c), with a significant decrease in synaptic puncta in hypoxia_500µM_ and an even more pronounced reduction in hypoxia_GD_ compared to hypoxia (p<0.01 and p<0.0001, respectively). In addition, hypoxia_GD_ exhibited a significant decrease compared to normoxia (p = 0.0023).

## 4. Discussion

In this study, we investigated human-derived *in vitro* model responses to different metabolic stress conditions in terms of electrophysiological activity and cell viability. The results showed that hypoxia alone, hypoxia_GD_, and hypoxia combined with different concentrations of glutamate yield distinct neuronal network responses over time.

### Oxygen-glucose deprivation

The results of the hypoxia_GD_ model revealed a distinct temporal pattern of neuronal network activity compared to hypoxia alone. Specifically, hypoxia alone induced a gradual and monotonic reduction in activity over time. In contrast, hypoxia_GD_ triggered a biphasic response, with a transient increase in activity during the first hours, followed by a rapid decline leading to network silencing. This early hyperactivation under hypoxia_GD_ is consistent with ATP depletion–induced disruption of ionic gradients and Na^+^/K^+^-ATPase failure, leading to membrane depolarization and increased release of excitatory neurotransmitters ^30–32^. However, this effect was short-lived and followed by a marked suppression of activity, which may reflect calcium overload and synaptic dysfunction ^33,34^. Hypoxia_GD_ has been shown to impair synaptic transmission primarily through impaired transmitter release^35^ followed by calcium-dependent mechanisms and alterations in receptor function^33^. These changes are reversible if oxygen and glucose delivery are restored in time ^35–37^, but persistent hypoxia_GD_ may lead to enduring network instability and collapse ^21,33,34^.

Despite these different temporal dynamics, both conditions converged toward severe functional impairment over 48 hours of metabolic stress and showed similarly limited functional recovery after reoxygenation. However, hypoxia_GD_ was associated with a higher percentage of dead cells and a stronger reduction in synaptic puncta compared to hypoxia. These findings are consistent with the well-established role of metabolic failure in driving oxidative stress, mitochondrial dysfunction, and apoptotic pathways leading to progressive neuronal damage previously observed in glucose deprivation models under experimental *in vitro* conditions ^21^. Our findings indicate that, while network-level activity may converge to a similar functional endpoint, the underlying structural neuronal damage is more severe under hypoxia_GD_. We assume that synaptic failure by presynaptic interruption of transmitter release is a key mechanism of neuronal network dysfunction in the absence of structural damage ^35^.

While the present findings provide a comprehensive characterization of neuronal network, sample size was relatively limited, which may contribute to variability in some of the reported effects.

### Hypoxia combined with glutamate

Glutamate exposure under hypoxia altered the temporal dynamics of network activity. Especially for the higher concentrations, a triphasic response was observed, with an initial reduction in activity followed by a transient increase and a subsequent decline, leading to network silencing. At high concentrations (500 µM glutamate), network bursting rapidly suppressed and completely abolished. These dynamics are consistent with excitotoxic mechanisms observed in human neuronal *in vitro* models^13,22,38,39^and *in vivo* animal models^40,41^, where impaired energy metabolism leads to extracellular glutamate accumulation, overstimulation of NMDA and AMPA receptors, and calcium overload, ultimately disrupting neuronal signalling ^13,22,38,39^. These are also present under hypoxia alone but are accelerated and amplified in the presence of exogenous glutamate^10,38,39^.

Neuronal activity patterns were different with lower glutamate concentrations. Networks exposed to 100 µM glutamate showed good functional recovery, with several parameters returning to levels not significantly different from baseline. The hypoxia_200µM_ condition showed partial recovery, whereas both hypoxia alone and hypoxia_500µM_ showed poor recovery, with similarly reduced activity levels. These results indicate that low glutamate exposure under hypoxia may allow partial restoration of network activity after reoxygenation. This is in line with previous work, where we showed that mild neuronal stimulation acts neuroprotective and improves neuronal network recovery during and after metabolic stress ^10,17,20,42,43^. We hypothesize that low concentrations of glutamate can be neuroprotective, especially in the case of synaptic arrest, while high glutamate concentrations are detrimental through excitotoxicity pathways. However, we did not assess cellular energy metabolism (e.g., ATP levels) or NMDA receptor activity, which could have provided additional mechanistic insight into the observed electrophysiological alterations.

Although functional outcomes were broadly similar across neuronal networks under hypoxia and hypoxia_500µM_, the cellular and structural analyses revealed important differences during the hypoxic phase. In particular, the hypoxia_500µM_ condition exhibited a higher proportion of live cells compared to hypoxia alone, yet this was accompanied by a significant reduction in synaptic puncta, indicating synaptic damage. This suggests that neuronal survival and synaptic integrity are differentially affected by glutamate exposure.

Mechanistically, dissociation between cell viability and synaptic structure is consistent with previously reported excitotoxic processes, where glutamate-induced NMDA receptor overactivation led to calcium overload and preferential damage to synaptic compartments ^44^. In this context, synaptic loss and functional disconnection may occur independently of, and potentially prior to, overt neuronal death ^44–46^.

## Conclusions

Our findings show that different forms of metabolic and excitotoxic stress produce distinct effects on neuronal networks in terms of temporal dynamics and cellular and synaptic alterations. Hypoxia_GD_ was associated with more cell death than hypoxia alone, whereas high concentrations of glutamate during hypoxia were associated with specific damage on the synaptic level. On the other hand, low concentrations of glutamate brought improved neuronal network activity, supporting previous results on neuroprotection by mild neuronal activation. The combination of functional and structural readouts provides a comprehensive understanding of neuronal network dynamics under metabolic stress.

## Supporting information

upplementary Material

## 5. Author declarations

## Availability of data

The datasets used and analysed during the current study are available from the corresponding author on reasonable request.

