## Supplementary material for "Distinct effects of different metabolic stress models on human-derived neuronal networks": upplementary Material

#### Supplementary Figures

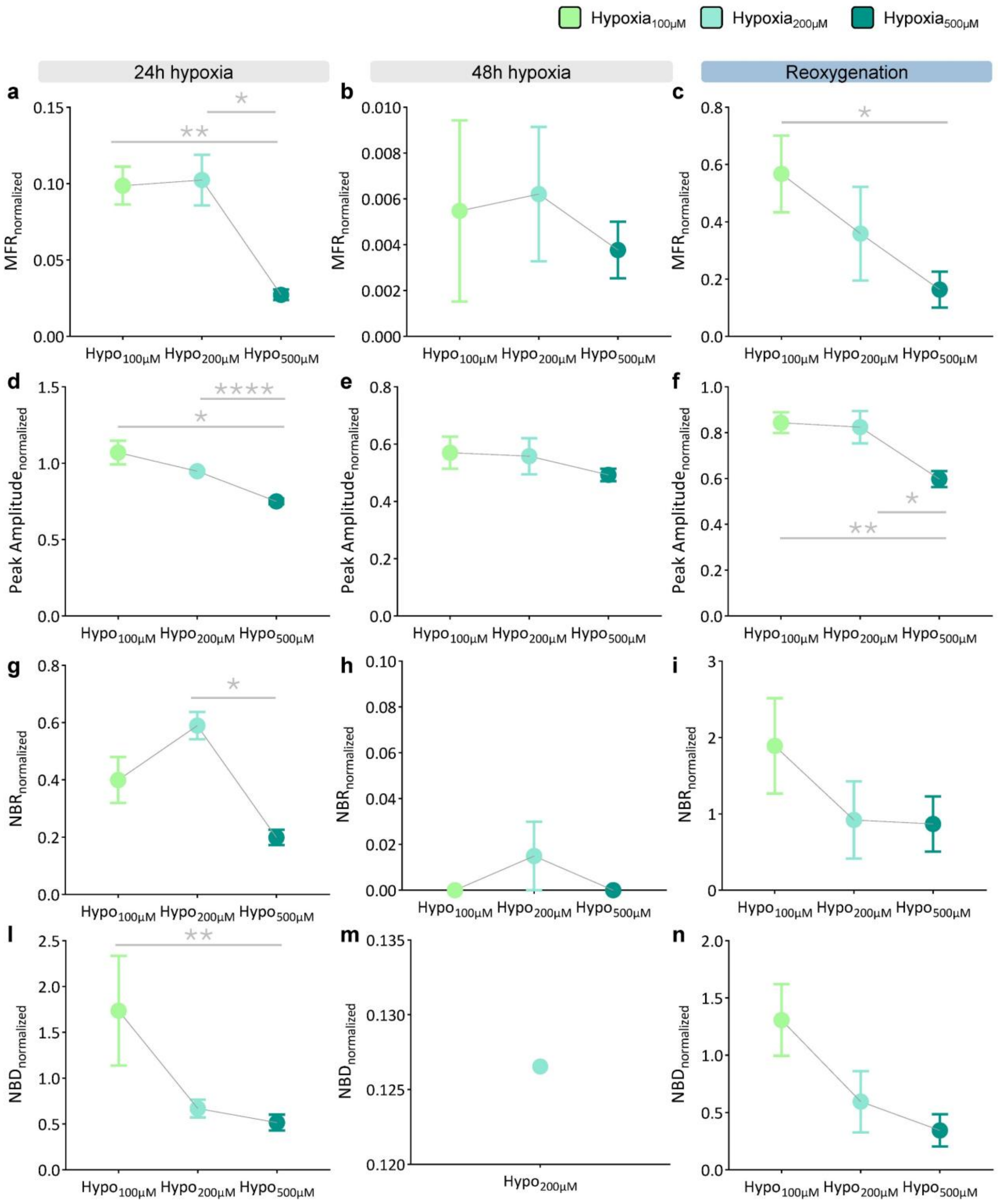

Figure S1. Curve dose response of glutamate. a-c) Graphs showing the normalized firing rate profile during: 24 hours of hypoxia (a); 48 hours of hypoxia (b); 24 hours of reoxygenation (c). d-f) Graphs showing normalized peak amplitude profile during: 24 hours of hypoxia (d); 48 hours of hypoxia (e); 24 hours of reoxygenation (f). g-i) Graphs showing normalized network bursting rate (d) rate profile during: 24 hours of hypoxia (g); 48 hours of hypoxia (h); 24 hours of reoxygenation (i). l-n) Graphs showing normalized network burst duration profile during: 24 hours of hypoxia (l); 48 hours of hypoxia (m); 24 hours of reoxygenation (n). Data are shown with the mean (dots) and standard error of the mean (whiskers). *p < 0.05, **p < 0.01, ***p < 0.001, ****p < 0.0001.

### Supplementary Tables

| | 2 hours | 6 hours | 12 hours | 18 hours | 24 hours | 30 hours | 36 hours | 42 hours | 48 hours |
|---|---|---|---|---|---|---|---|---|---|
| $Hypo_{0\mu M}$-$Hypo_{100\mu M}$ | 0.023829 | 0.025725 | ns | ns | ns | ns | ns | ns | ns |
| $Hypo_{0\mu M}$-$Hypo_{200\mu M}$ | 0.000098 | 0.000047 | 0.000474 | ns | ns | ns | ns | ns | ns |
| $Hypo_{0\mu M}$-$Hypo_{500\mu M}$ | 0.000280 | 0.000183 | 0.000193 | 0.006238 | 0.034996 | ns | ns | ns | ns |
| $Hypo_{100\mu M}$-$Hypo_{200\mu M}$ | 0.003784 | 0.008541 | 0.029334 | ns | ns | ns | ns | ns | ns |
| $Hypo_{100\mu M}$-$Hypo_{500\mu M}$ | 0.006526 | 0.008842 | 0.002787 | 0.0000440 | 0.012876 | 0.026833 | ns | ns | ns |
| $Hypo_{200\mu M}$-$Hypo_{500\mu M}$ | ns | ns | 0.028774 | 0.003607 | 0.030047 | ns | ns | ns | ns |

Table S1: Statistical analysis results of the mean firing rate comparing the $hypoxia_{0\mu M}$, $hypoxia_{100\mu M}$, $hypoxia_{200\mu M}$, and $hypoxia_{500\mu M}$ at the different time point during the 48 hours of hypoxia.

| | 2 hours | 6 hours | 12 hours | 18 hours | 24 hours | 30 hours | 36 hours | 42 hours | 48 hours |
|---|---|---|---|---|---|---|---|---|---|
| $Hypo_{0\mu M}$-$Hypo_{100\mu M}$ | ns | ns | ns | ns | ns | ns | ns | ns | ns |
| $Hypo_{0\mu M}$-$Hypo_{200\mu M}$ | ns | 0.013902 | ns | ns | ns | ns | ns | ns | ns |
| $Hypo_{0\mu M}$-$Hypo_{500\mu M}$ | ns | ns | ns | ns | 0.000269 | 0.001129 | ns | ns | ns |
| $Hypo_{100\mu M}$-$Hypo_{200\mu M}$ | ns | 0.038737 | ns | ns | ns | ns | ns | ns | ns |
| $Hypo_{100\mu M}$-$Hypo_{500\mu M}$ | ns | ns | ns | ns | ns | 0.040957 | ns | ns | ns |
| $Hypo_{200\mu M}$-$Hypo_{500\mu M}$ | ns | ns | ns | ns | 0.001298 | 0.015463 | ns | ns | ns |

Table S2: Statistical analysis results of the peak amplitude comparing the $hypoxia_{0\mu M}$, $hypoxia_{100\mu M}$, $hypoxia_{200\mu M}$, and $hypoxia_{500\mu M}$ at the different time point during the 48 hours of hypoxia.

| | 2 hours | 6 hours | 12 hours | 18 hours | 24 hours | 30 hours | 36 hours | 42 hours | 48 hours |
|---|---|---|---|---|---|---|---|---|---|
| $Hypo_{0\mu M}$-$Hypo_{100\mu M}$ | 0.012759 | ns | ns | ns | ns | ns | ns | ns | ns |
| $Hypo_{0\mu M}$-$Hypo_{200\mu M}$ | 0.000921 | 0.004076 | ns | ns | ns | 0.014239 | ns | ns | ns |
| $Hypo_{0\mu M}$-$Hypo_{500\mu M}$ | 0.000219 | 0.000054 | 0.000087 | ns | ns | ns | ns | ns | ns |
| $Hypo_{100\mu M}$-$Hypo_{200\mu M}$ | ns | ns | ns | ns | ns | ns | ns | ns | ns |
| $Hypo_{100\mu M}$-$Hypo_{500\mu M}$ | 0.004146 | 0.002969 | 0.005423 | 0.014751 | ns | ns | ns | ns | ns |
| $Hypo_{200\mu M}$-$Hypo_{500\mu M}$ | ns | 0.028892 | 0.000163 | 0.000267 | 0.000884 | 0.009502 | ns | ns | ns |

Table S3: Statistical analysis results of the network bursting rate comparing the $hypoxia_{0\mu M}$, $hypoxia_{100\mu M}$, $hypoxia_{200\mu M}$, and $hypoxia_{500\mu M}$ at the different time point during the 48 hours of hypoxia.

| | 2 hours | 6 hours | 12 hours | 18 hours | 24 hours | 30 hours | 36 hours | 42 hours | 48 hours |
|---|---|---|---|---|---|---|---|---|---|
| $Hypo_{0µM}$-$Hypo_{100µM}$ | ns | ns | ns | ns | ns | ns | ns | ns | ns |
| $Hypo_{0µM}$-$Hypo_{200µM}$ | 0.000038 | 0,000036 | 0,000430 | ns | ns | ns | ns | ns | ns |
| $Hypo_{0µM}$-$Hypo_{500µM}$ | 0.000159 | 0,001716 | 0,000028 | 0,038218 | ns | ns | ns | ns | ns |
| $Hypo_{100µM}$-$Hypo_{200µM}$ | 0.000412 | 0,000079 | 0,002466 | ns | ns | 0.043317 | ns | ns | ns |
| $Hypo_{100µM}$-$Hypo_{500µM}$ | 0.000195 | 0,002476 | 0,000226 | 0,030782 | ns | ns | ns | <0,000000000000001 | ns |
| $Hypo_{200µM}$-$Hypo_{500µM}$ | ns | ns | 0,000300 | ns | ns | ns | ns | <0,000000000000001 | ns |

Table S4: Statistical analysis results of the network burst duration comparing the $hypoxia_{0µM}$, $hypoxia_{100µM}$, $hypoxia_{200µM}$, and $hypoxia_{500µM}$ at the different time point during the 48 hours of hypoxia.
